# Disrupted sleep in dystonia depends on cerebellar function but not motor symptoms in mice

**DOI:** 10.1101/2023.02.09.527916

**Authors:** Luis E. Salazar Leon, Roy V. Sillitoe

**Affiliations:** Department of Neuroscience, Baylor College of Medicine, Houston, Texas, USA; Department of Pathology & Immunology, Baylor College of Medicine, Houston, Texas, USA; Department of Pediatrics, Baylor College of Medicine, Houston, Texas, USA; Development, Disease Models & Therapeutics Graduate Program, Baylor College of Medicine, Houston, Texas, USA; Jan and Dan Duncan Neurological Research Institute at Texas Children’s Hospital, Houston, Texas, 77030, USA

**Keywords:** Dystonia, sleep, circadian rhythms, Purkinje cells, cerebellar nuclei

## Abstract

Although dystonia is the third most common movement disorder, patients often also experience debilitating nonmotor defects including impaired sleep. The cerebellum is a central component of a “dystonia network” that plays various roles in sleep regulation. Importantly, the primary driver of sleep impairments in dystonia remains poorly understood. The cerebellum, along with other nodes in the motor circuit, could disrupt sleep. However, it is unclear how the cerebellum might alter sleep and mobility. To disentangle the impact of cerebellar dysfunction on motion and sleep, we generated two mouse genetic models of dystonia that have overlapping cerebellar circuit miswiring but show differing motor phenotype severity: *Ptf1a*^*Cre*^*;Vglut2*^*fx/fx*^ and *Pdx1*^*Cre*^*;Vglut2*^*fx/fx*^ mice. In both models, excitatory climbing fiber to Purkinje cell neurotransmission is blocked, but only the *Ptf1a*^*Cre*^*;Vglut2*^*fx/fx*^ mice have severe twisting. Using *in vivo* ECoG and EMG recordings we found that both mutants spend greater time awake and in NREM sleep at the expense of REM sleep. The increase in awake time is driven by longer awake bouts rather than an increase in bout number. We also found a longer latency to reach REM in both mutants, which is similar to what is reported in human dystonia. We uncovered independent but parallel roles for cerebellar circuit dysfunction and motor defects in promoting sleep quality versus posture impairments in dystonia.

## Introduction

Dystonia presents with phenotypic and etiologic heterogeneity. Considered the third most common movement disorder, “dystonia” does not comprise a single disease or symptom, but rather describes an array of disorders sharing overlapping behavioral outcomes. While different forms of dystonia express unique etiologies, substantial evidence implicates the cerebellum as a major node in the underlying network disruptions^1–4^. Dysfunction of Purkinje cells and the cerebellar nuclei, the primary outputs of the cerebellar cortex and cerebellum, respectively, are implicated in both hereditary and idiopathic forms of dystonia^4–7^. Importantly, while cerebellar dysfunction is sufficient to induce dystonia in animal models, therapies addressing cerebellar dysfunction can modulate dystonia and reduce motor symptom severity. One such therapy, cerebellar deep brain stimulation (DBS), has been used to effectively reduce motor symptom severity in both mouse models^4^ and human patients^8,9^, further suggesting a critical cerebellar involvement in the etiology of dystonia. However, nonmotor behaviors are also relevant to the cerebellum and to dystonia.

Along with its known role in regulating motor function, increasing evidence shows that the cerebellum also serves as a key brain region in the control of a variety of nonmotor behaviors such as cognitive and emotional processing^10^, associative learning^11^, and reward expectation^11,12^. Recent work also suggests that the cerebellum may play a role in sleep-related behaviors. Purkinje cells and cerebellar nuclei neurons have been found to display sleep-dependent activity, increasing their firing during NREM (non-rapid eye movement) sleep^13–16^. In addition, lesioning of the cerebellar vermis has been shown to impair sleep, suggesting that normal cerebellar circuitry and activity are important for maintaining sleep rhythms^17,18^. Numerous studies have shown that cerebellar disruptions form the basis for the many comorbidities of motor disorders^19–22^. However, the role of the cerebellum and its circuit components in sleep regulation have not been studied thoroughly in dystonia. It is known that dystonic patients demonstrate increased sleep latency and REM (rapid eye movement) latency, and in some cases, persistent involuntary muscle contractions during sleep^23–25^. It is also noted that therapeutics which successfully alleviate the motor symptoms of dystonia appear to have little to no effect on the sleep disruptions^26^. The importance of addressing sleep dysfunction is becoming increasingly apparent in society, as sleep disruptions can significantly impact quality of life, driving many subsequent comorbidities^27,28^. Disrupted sleep is also associated with impaired motor learning/function^29–31^, as synaptic activity is normalized during sleep^32^. Together, the mounting evidence inspires a compelling model in which sleep and motor dysfunction in dystonia comprise two halves of a self-propelling cycle. It is possible that cerebellar dysfunction drives both the more commonly-appreciated motor abnormalities and the nonmotor sleep disruptions, although how they emerge needs to be systematically resolved.

It remains unclear whether dystonic motor dysfunctions persist during all stages of sleep, and consistently across different manifestations of the disease; recent evidence from a survey of cervical dystonia patients suggests that, in cases of idiopathic cervical dystonia, it does^24^. The lack of clarity on which factors (motor dysfunction and/or cerebellar dysfunction) drive sleep dysfunction in dystonia highlights the significance of this knowledge gap. To investigate the relationship between cerebellar dysfunction, motor dysfunction, and sleep, here we used a constitutively active Cre/*lox*-p system to drive the deletion of the *Vglut2* gene in afferent neurons that project excitatory fibers that ultimately communicate with the Purkinje cells. *Vglut2* was deleted using the *Ptf1a* and *Pdx1* gene regulatory elements to spatially drive Cre expression: the resulting mice had the genotypes *Ptf1a*^*Cre*^*;Vglut2*^*fx/fx*^ and *Pdx1*^*Cre*^*;Vglut2*^*fx/fx*^. The *Ptf1a* and *Pdx1* genes are expressed in the excitatory neurons of the inferior olive, a region of the brainstem, which projects afferent fibers to the cerebellar cortex and terminate as excitatory climbing fibers^4,33^. However, *Ptf1a* expression occurs in a wider distribution across the inferior olive relative to *Pdx1*, and *Pdx1* is also expressed in mossy fiber afferent neurons (Lackey et al., 2023 *in preparation*). Therefore, the silencing of excitatory climbing fiber synapses occurs with differential coverage in mice with *Ptf1a-* versus *Pdx1*-driven Cre expression. Only *Ptf1a*^*Cre*^*;Vglut2*^*fx/fx*^ mice present with severe motor dysfunction involving twisting of the torso and hyperextensions of the back and limbs^4^, while *Pdx1*^*Cre*^*;Vglut2*^*fx/fx*^ adult mice show only subtle dystonic behaviors^34^. Thus, the use of both mouse models allows us to deliberately silence excitatory olivocerebellar synapses while also providing us with an opportunity to query varying severities of cerebellar dysfunction.

In this work, we report that, despite their different dystonia-related motor severities, *Pdx1*^*Cre*^*;Vglut2*^*fx/fx*^ and *Ptf1a*^*Cre*^*;Vglut2*^*fx/fx*^ mice display similarly impaired sleep physiology and circadian rhythms. Both mutants display highly disrupted sleep, spending greater time awake and at the expense of REM sleep. Furthermore, we found that both the *Pdx1*^*Cre*^*;Vglut2*^*fx/fx*^ and *Ptf1a*^*Cre*^*;Vglut2*^*fx/fx*^ mutant mice display increased latency to reach REM, which is similar to what is observed in human patients with dystonia^23^. Intriguingly, only mice with overt dystonic motor behaviors (*Pt1fa*^*Cre*^*;Vglut2*^*fx/fx*^) show differences in ECoG spectral power frequency, particularly in the latter half of the time spent asleep. We also found that circadian activity rhythms remain unchanged across all groups of mice, and that the circadian “master clock” remains ostensibly unaffected by our circuit manipulation. Our work demonstrates that aberrant cerebellar activity dually disrupts motor function and sleep, paving the way for improved future therapeutics that may be able to simultaneously address both motor and sleep dysfunction in the context of motor disease.

## Results

### *Ptf1a*^*Cre*^*;Vglut2*^*fx/fx*^ and *Pdx1*^*Cre*^*;Vglut2*^*fx/fx*^ mice display overlapping cerebellar circuit deficits, but only the *Ptf1a*^*Cre*^*;Vglut2*^*fx/fx*^ mice show overt dystonic symptoms

We have previously demonstrated that silencing glutamatergic olivocerebellar synapses can induce severe dystonic motor phenotypes^4^. To elucidate the relative contributions of cerebellar versus motor dysfunction on sleep impairments, we additionally leveraged a previously generated mouse model of cerebellar dysfunction lacking overt motor dysfunction: the *Pdx1*^*Cre*^*;Vglut2*^*fx/fx*^ mouse^34^ (Supplementary video 1). Both the *Ptf1a*^*Cre*^*;Vglut2*^*fx/fx*^ and *Pdx1*^*Cre*^*;Vglut2*^*fx/fx*^ mouse models utilize the Cre/*lox*-p system to drive the deletion of *Vglut2*, but under different promoters (Figure 1A). A detailed characterization of the *Ptf1a*^*Cre*^*;Vglut2*^*fx/fx*^ mice and their resulting dystonia were previously described^4^, whereas the behavior and circuit basis of the *Pdx1*^*Cre*^*;Vglut2*^*fx/fx*^ mice will be described extensively in an independent study (Lackey et al., 2023 *in preparation*). While both models result in the loss of VGLUT2 protein after genetically targeting glutamatergic olivocerebellar synapses in Purkinje cells (Figure 1B-C), the differential expression patterns of *Ptf1a* (in climbing fiber neurons) and *Pdx1* (in climbing fiber and mossy fiber neurons) yield differential synapse silencing resulting in different motor phenotypes (Figure 1D, Figure 1 supplement 1). Compared to wildtype littermate controls (Figure 1E), the *Ptf1a*^*Cre*^*;Vglut2*^*fx/fx*^ mice display overt dystonic motor phenotypes and dystonic behaviors (Figure 1F), while the *Pdx1*^*Cre*^*;Vglut2*^*fx/fx*^ mice do not display overt dystonic motor phenotypes such as spontaneous twisting postures and hyperextension of the back, limbs, or digits (Figure 1G). To better understand the alterations in motor phenotypes in both mouse models, we implanted mice with cortical (ECoG) and muscular (EMG) electrodes for *in vivo* monitoring. We calculated the overall EMG power in the 0-30Hz frequency range, which has been previously used to quantitatively diagnose dystonia in human patients^35^ (representative EMG traces per genotype; Figure 1H). As predicted, the overall EMG power was significantly elevated in the *Ptf1a*^*Cre*^*;Vglut2*^*fx/fx*^ mice as compared to the control and *Pdx1*^*Cre*^*;Vglut2*^*fx/fx*^ mice (Figure 1H-I). The EMG activity reflects prolonged over-contractions, which is a key phenotype observed in human patients with generalized and focal dystonia. Additionally, we found that this elevation of cervical EMG activity was maintained during all states, including both REM and NREM sleep (Figure 1 supplement 2, B-E). These findings support the idea that *Ptf1a*^*Cre*^*;Vglut2*^*fx/fx*^ mice display elevated muscle activity due to cerebellar dysfunction, while *Pdx1*^*Cre*^*;Vglut2*^*fx/fx*^ mice do not. These results further suggest that cerebellar circuit manipulations can occur without causing overt and severe motor dysfunctions and furthermore establishes the two mouse models for use in subsequent experiments in this study.

**Figure 1:**
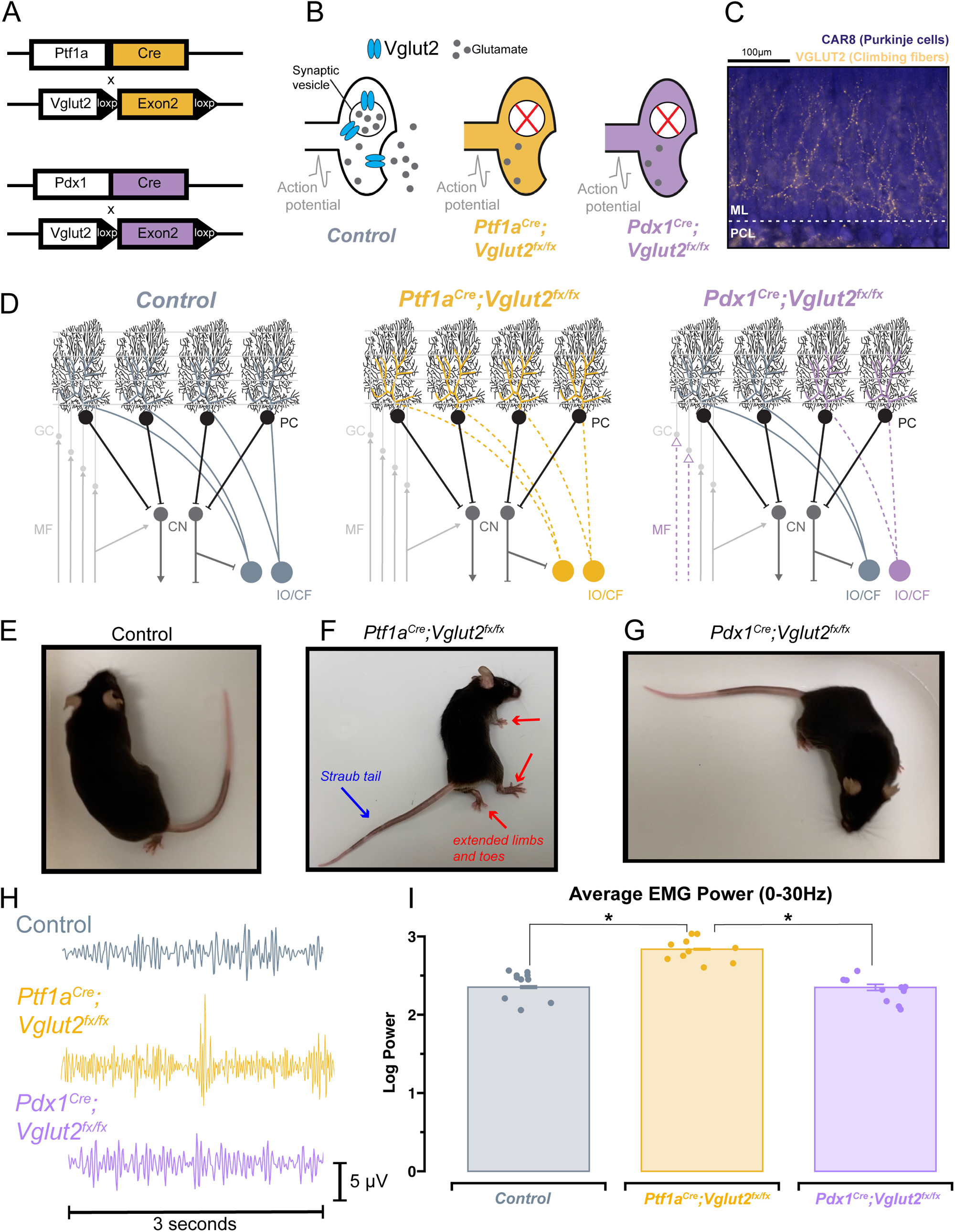
*Ptf1a*^*Cre*^*;Vglut2*^*fx/fx*^ and *Pdx1*^*Cre*^*;Vglut2*^*fx/fx*^ mice display differential dystonic motor phenotypes. **(a)** Using either the *Ptf1a*^*Cre*^ or *Pdx1*^*Cre*^ genetic driver lines, exon 2 of *Vglut2* was selectively removed and as a result VGLUT2 expression was deleted with spatial specificity. **(b)** Schematic illustration demonstrating the result of the *Vglut2* deletion and the subsequent synaptic silencing in the affected cells. **(c)** Immunohistochemical staining of the cerebellar cortex, showing Purkinje cells (blue) and VGLUT2-positive climbing fibers from the inferior olive (gold). Abbreviations: ML, molecular layer; PCL, Purkinje cell layer. **(d)** Schematic demonstrating the end-result of the *Vglut2* deletion in the *Ptf1a*^*Cre*^*;Vglut2*^*fx/fx*^ and *Pdx1*^*Cre*^*;Vglut2*^*fx/fx*^ mice. *Ptf1a*^*Cre*^*;Vglut2*^*fx/fx*^ mice have widespread silencing of olivocerebellar glutamatergic synapses, while *Pdx1*^*Cre*^*;Vglut2*^*fx/fx*^ mice have comparatively more restricted silencing of a subset of olivocerebellar synapses. Abbreviations: GC, granule cell; MF, mossy fiber; PC, Purkinje cell, CF, climbing fiber; CN, cerebellar nuclei; IO, inferior olive. **(e)** Video still from a control mouse with no atypical function. **(f)** Video still demonstrating dystonic postures in a *Ptf1a*^*Cre*^*;Vglut2*^*fx/fx*^ mouse, specifically showing the hindlimb hyperextension and Straub tail (noted by red and blue arrows). **(g)** Video still from a *Pdx1*^*Cre*^*;Vglut2*^*fx/fx*^ mouse demonstrating the absence of overt dystonic motor dysfunction. **(h)** Raw EMG waveforms of trapezius muscle activity for a 3-second period. Control (grey), *Ptf1a*^*Cre*^*;Vglut2*^*fx/fx*^ (gold), and *Pdx1*^*Cre*^*;Vglut2*^*fx/fx*^ (purple) mice. **(i)** Quantification of the overall EMG activity (0-30Hz) for all mice used in the study. Points on **i** represent individual mice, n=10 per group. Source data and specific p-values for **i** are available in Figure 1-source data 1.

### Circadian activity is unchanged in *Ptf1a*^*Cre*^*;Vglut2*^*fx/fx*^ and *Pdx1*^*Cre*^*;Vglut2*^*fx/fx*^ mutant mice

Wheel-running activity is commonly used in rodents as a proxy for measuring daily activity patterns^36^. Previous work has used wheel-running to understand circadian activity in other mouse models of movement disorders (for example, mild ataxia)^37^. Therefore, we sought to determine the extent of circadian activity disruption in *Ptf1a*^*Cre*^*;Vglut2*^*fx/fx*^ and *Pdx1*^*Cre*^*;Vglut2*^*fx/fx*^ mutant mice. Mice were singly housed with *ad libitum* access to food, water, and a running wheel in their home cage (Figure 2A). Wheel revolutions were automatically monitored throughout the recording period that lasted 35-days (14 days baseline (LD; light-dark), 21 days constant condition (DD; dark-dark periods; Figure 2B)). The collected data were analyzed and plotted as actograms for ease of viewing; each row represents a day and black tick marks represent revolutions of the running wheel, indicative of locomotor activity. Data is double plotted (as convention), such that 48-hours of activity are plotted on the same line, to better visualize changes in activity patterns^36^. We predicted that differences in circadian activity patterns in our mutant mice would arise either from motor dysfunction or our cerebellar circuit manipulation. Given that *Pdx1*^*Cre*^*;Vglut2*^*fx/fx*^ mice do not display severe dystonic motor behaviors, and the extent of their olivocerebellar manipulation is restricted relative to *Ptf1a*^*Cre*^*;Vglut2*^*fx/fx*^ mice, we predicted that their circadian activity profiles would remain unchanged relative to littermate controls. As expected, we observed normal wheel-running behavior in *Pdx1*^*Cre*^*;Vglut2*^*fx/fx*^ mice relative to littermate controls (Figure 2C/E). We also found that despite their overt motor dysfunction, *Ptf1a*^*Cre*^*;Vglut2*^*fx/fx*^ mice did voluntarily run on the wheel and maintain rhythmic behavior, though after a longer (8-days) acclimation period (Figure 2D). We observed significantly lower average activity counts for *Ptf1a*^*Cre*^*;Vglut2*^*fx/fx*^ mice with overt motor dysfunction during both LD and DD paradigms (Figure 2F-G). Nevertheless, both *Pdx1*^*Cre*^*;Vglut2*^*fx/fx*^ and *Ptf1a*^*Cre*^*;Vglut2*^*fx/fx*^ mice displayed characteristic nocturnal behavior even during the DD phase, similar to controls (Figure 2C-E). We also assessed endogenous circadian period length (tau), a measure of the period of a circadian rhythm. The tau length refers to the length of time it takes for the rhythm to complete one cycle^36^. At the end of the DD paradigm, all mice displayed an average tau of ~23.7hrs (Figure 2H). This slight deviation from 24hrs is expected, as endogenous tau length in mice is slightly less than 24hrs^36^. In addition, we also found that the “siesta” period — a brief bout of sleep during the active period^38^ — in *Ptf1a*^*Cre*^*;Vglut2*^*fx/fx*^ mice is significantly longer by 7-10 minutes (Figure 2I). However, this increase may be a result of the low background activity for *Ptf1a*^*Cre*^*;Vglut2*^*fx/fx*^ mice. To validate that our genetic manipulation of *Vglut2* did not significantly alter the major sleep center of the brain, we assessed *Vglut2* mRNA expression in control *Ptf1a*^*Cre*^ and *Pdx1*^*Cre*^ mice (without *floxed* alleles of *Vglut2*) using *in situ* hybridization. We found a lack of *Vglut2* expression in the suprachiasmatic nucleus (SCN) “master clock”. This was anticipated since the SCN is a heavily GABAergic region^39^ (*Vgat*-expressing, Figure 2 supplement 1). These results suggest that in the *Pdx1*^*Cre*^*;Vglut2*^*fx/fx*^ and *Ptf1a*^*Cre*^*;Vglut2*^*fx/fx*^ mice, circadian rhythms remain largely unchanged despite cerebellar and motor dysfunction.

**Figure 2:**
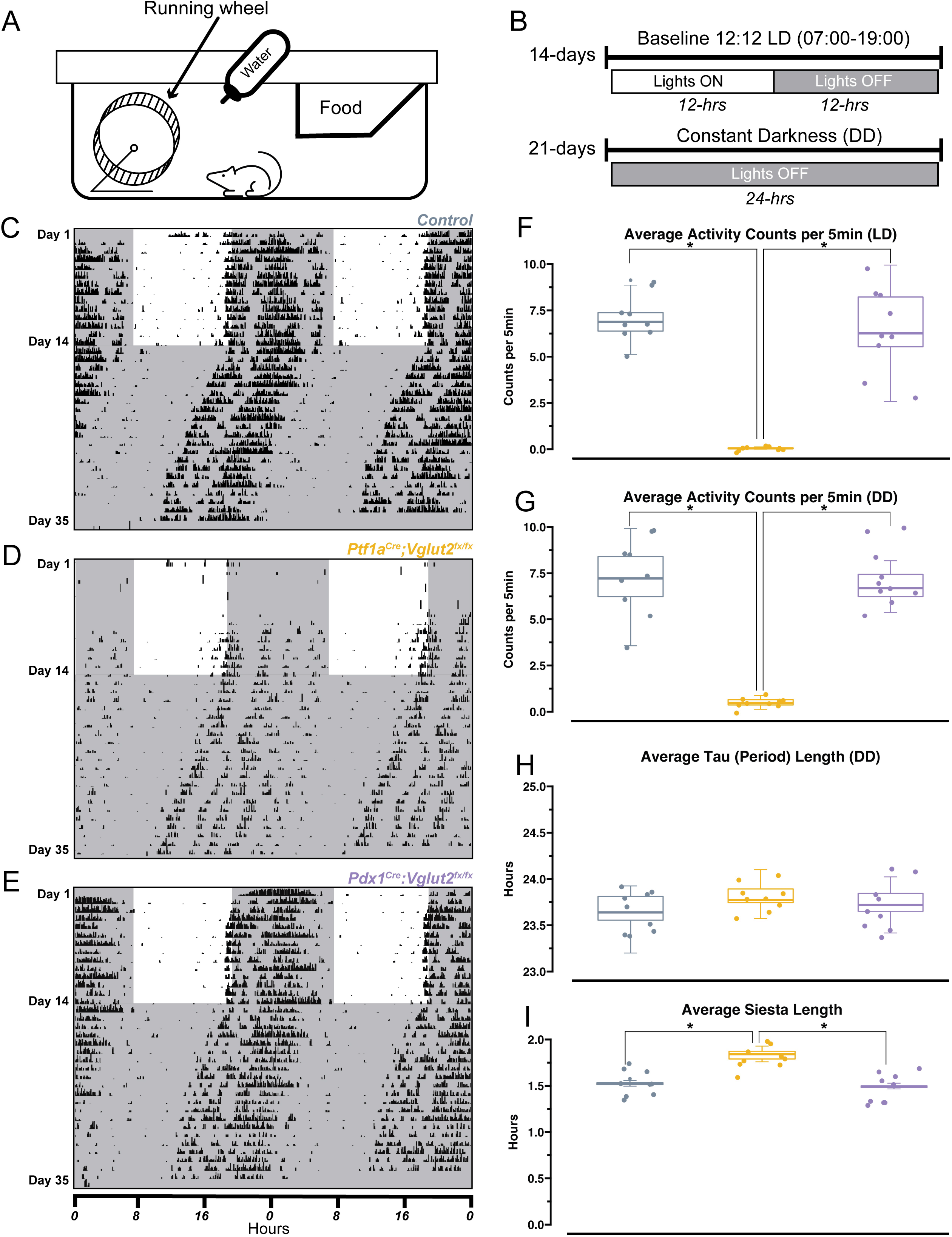
*Ptf1a*^*Cre*^*;Vglut2*^*fx/fx*^ and *Pdx1*^*Cre*^*;Vglut2*^*fx/fx*^ mice display normal circadian rhythmicity. **(a)** Schematic illustration of the wheel-running setup. **(b)** Timeline of wheel-running experiment. **(c)** Representative double-plotted actogram for a control mouse. Each row represents a day, black tick marks represent wheel-running activity (measured as revolutions of the running wheel). Black shaded regions represent “lights OFF”, unshaded regions represent “lights ON”. **(d)** Same as **(c)** but for a *Pdx1*^*Cre*^*;Vglut2*^*fx/fx*^ mouse. **(e)** Same as **(c)** but for a *Ptf1a*^*Cre*^*;Vglut2*^*fx/fx*^ mouse. **(f)** Quantification of average activity counts per 5min for all mice, only during the LD paradigm. **(g)** Same as **(f)** but only quantifying activity during the DD paradigm. **(h)** Quantification of circadian period (tau) for all mice, during the DD paradigm. **(i)** Quantification of “siesta” bout length for all mice throughout the 35-day recording period. Points on **f-i** represent individual mice, n=9 mice per group. The source data and specific p-values for **f-i** are available in Figure 2-source data 1.

### Cerebellar dysfunction disrupts sleep stages independently of the dystonic phenotype

The relationship between sleep and motor function is particularly relevant in dystonia, as reports suggest that motor symptoms are easier to manage after a good night’s sleep, and earlier in the morning, shortly after waking up^40^. Therefore, a major goal was to determine the overall sleep quality in *Ptf1a*^*Cre*^*;Vglut2*^*fx/fx*^ and *Pdx1*^*Cre*^*;Vglut2*^*fx/fx*^ mice. We implanted *Ptf1a*^*Cre*^*;Vglut2*^*fx/fx*^ and *Pdx1*^*Cre*^*;Vglut2*^*fx/fx*^ mice with ECoG/EMG electrodes that were made out of silver wire and recorded signals continuously for 8-hrs during the light phase (Figure 3A-C). Raw ECoG/EMG waveforms show that both *Ptf1a*^*Cre*^*;Vglut2*^*fx/fx*^ and *Pdx1*^*Cre*^*;Vglut2*^*fx/fx*^ mice display characteristic spectral activity which defines wake, NREM, and REM sleep (Figure 3D). We also note that high-amplitude spikes in the EMG activity were observed in *Ptf1a*^*Cre*^*;Vglut2*^*fx/fx*^ mice during brief periods of wake, indicative of motor dysfunction, while no such phenomenon was observed in the *Pdx1*^*Cre*^*;Vglut2*^*fx/fx*^ mice (Figure 3D). We then assessed the total time spent awake, in NREM sleep, and in REM sleep. While sleep cycles in mice are shorter than in humans, they do follow the similar pattern of wake, followed by NREM, and then REM sleep (Figure 3E). Representative hypnograms of 1-hour of total recording time showed that both the *Ptf1a*^*Cre*^*;Vglut2*^*fx/fx*^ and *Pdx1*^*Cre*^*;Vglut2*^*fx/fx*^ mice displayed disrupted sleep. The periods of wake were more frequent and last longer compared to the littermate controls (Figure 3F). We found that both *Pdx1*^*Cre*^*;Vglut2*^*fx/fx*^ and *Ptf1a*^*Cre*^*;Vglut2*^*fx/fx*^ mice spent more time awake and in NREM at the expense of decreased REM sleep (Figure 3G-I). These results suggest that, although motor dysfunction may occur in brief periods of spontaneous wakefulness (*Ptf1a*^*Cre*^*;Vglut2*^*fx/fx*^), cerebellar dysfunction alone may be sufficient to alter sleep activity independent from gross motor dysfunction (*Pdx1*^*Cre*^*;Vglut2*^*fx/fx*^).

**Figure 3:**
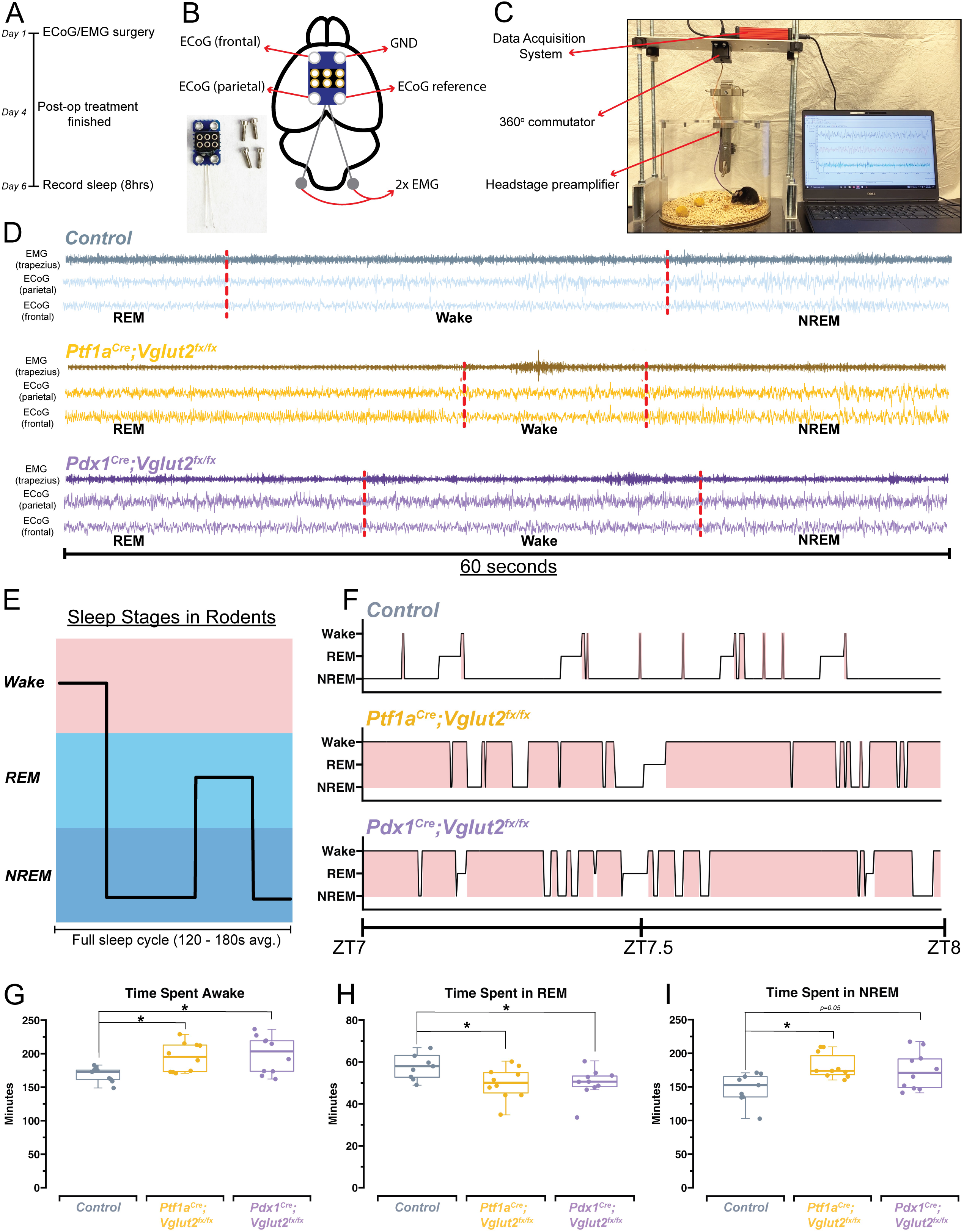
*Ptf1a*^*Cre*^*;Vglut2*^*fx/fx*^ and *Pdx1*^*Cre*^*;Vglut2*^*fx/fx*^ mice display disrupted sleep patterns. **(a)** Schematic illustration of the experimental timeline. **(b)** Schematic illustration of a mouse brain showing the placement of the ECoG/EMG headmount. An image of the headmount and mounting screws is also shown in the bottom left. **(c)** Video still from a sample sleep recording showing the experimental setup while a mouse is being recorded. **(d)** Raw waveforms of EMG activity (top trace for each sample) and ECoG activity (bottom two traces for each sample) for representative mice from each group. Each example is 60-seconds in length. Sleep stage, as determined by SPINDLE (see Methods), is noted under each example. Dotted red lines are added to help distinguish each sleep/wake state. **(e)** Schematic showing sleep stages and their organization for a mouse. **(f)** Hypnograms for a single representative mouse from each group, for the same 1-hr period, ZT7-ZT8, where ZT0 = lights ON, ZT1 = 1-hr after lights ON, etc. Periods of wake are highlighted in red. **(g)** Quantification of total time spent awake for all mice in each group. **(h)** Quantification of the total time spent in REM for all mice in each group. **(i)** Quantification of the total time spent in NREM for all mice in each group. Points on **g-i** represent individual mice, n=10 mice per group. The source data and specific p-values for **g-i** are shown in Figure 3-source data 1.

### Disrupted sleep patterns occur independent of overt dystonic motor dysfunction when comparing *Ptf1a*^*Cre*^*;Vglut2*^*fx/fx*^ and *Pdx1*^*Cre*^*;Vglut2*^*fx/fx*^ models of dystonia

We observed that cerebellar dysfunction was sufficient to disrupt sleep stages in *Ptf1a*^*Cre*^*;Vglut2*^*fx/fx*^ and *Pdx1*^*Cre*^*;Vglut2*^*fx/fx*^ mice, mouse models with and without overt dystonic motor phenotypes, respectively. However, the fluctuations in the frequency of sleep stages or length of sleep states, both of which could be driving the observed differences in sleep versus awake time, remained unclear (Figure 4A). Therefore, to further understand the fragility of sleep stages, and the disruption of each stage, we calculated both the total number of sleep-stage bouts along with the average length of bouts for wake, NREM, and REM. We note that these calculations were performed after the onset of sleep, which was determined using a similar approach to previous work^41^ (Figure 4B). We found that the total number of wake bouts was not different between *Ptf1a*^*Cre*^*;Vglut2*^*fx/fx*^, *Pdx1*^*Cre*^*;Vglut2*^*fx/fx*^, and the littermate controls (Figure 4C). However, for both *Pdx1*^*Cre*^*;Vglut2*^*fx/fx*^ and *Ptf1a*^*Cre*^*;Vglut2*^*fx/fx*^ mutant mice, the awake bouts were significantly longer than in controls, by an average of ~67 minutes (Figure 4D). To examine the disruptions in sleep stages after sleep onset, we calculated the total number of REM and NREM bouts. We found that both the *Pdx1*^*Cre*^*;Vglut2*^*fx/fx*^ and *Ptf1a*^*Cre*^*;Vglut2*^*fx/fx*^ mice displayed an increase in the overall number of NREM bouts coupled with fewer REM bouts (Figure 4G, 4E), while the average length of both REM and NREM bouts remained the same between all groups (Figure 4H, 4F). Previous work in human patients with cervical dystonia suggests that dystonic patients display an increased latency to sleep, with a particular effect on the REM stage of sleep^23^. We hypothesized that these phenotypes may be recapitulated in the *Pdx1*^*Cre*^*;Vglut2*^*fx/fx*^ and *Ptf1a*^*Cre*^*;Vglut2*^*fx/fx*^ mice. Therefore, we calculated the latency to reach REM and NREM sleep, as this could further indicate whether sleep dysfunction is primarily related to falling asleep versus staying asleep (or both). While both groups of mutant mice displayed a normal latency to reach NREM sleep (Figure 4J), latency to reach REM sleep was significantly elevated in both groups by an average of 47 minutes (Figure 4I). Together, these experiments highlight the specific deficits of sleep architecture that have been disordered in the *Pdx1*^*Cre*^*;Vglut2*^*fx/fx*^ and *Ptf1a*^*Cre*^*;Vglut2*^*fx/fx*^ mutant mice, and that the same deficits occur in both groups independently of how severe and constant the motor phenotype may be.

**Figure 4:**
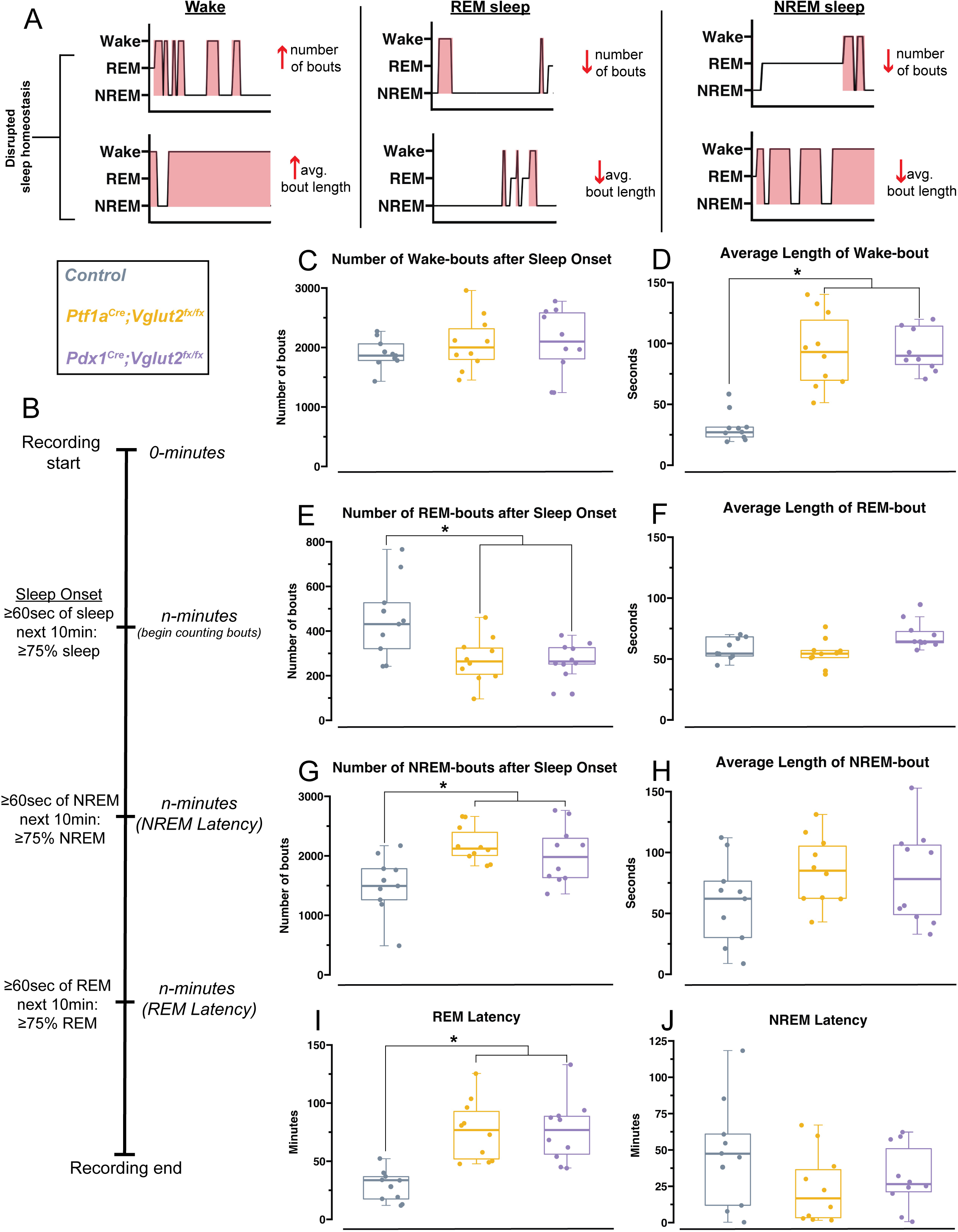
*Pdx1*^*Cre*^*;Vglut2*^*fx/fx*^ and *Ptf1a*^*Cre*^*;Vglut2*^*fx/fx*^ mutant mice display equivalent impairments in sleep. Schematic with hypnogram examples of how possible forms of sleep disruption may appear. Schematic showing sleep recording timeline and how sleep onset, NREM latency, and REM latency were defined and calculated, similar to Hunsley & Palmiter, 2004^41^. **(c)** Quantification of the total number of wake bouts. **(d)** Quantification of the average length of wake bouts. **(e)** Quantification for the number of REM bouts after sleep onset. **(f)** Quantification for the average length of REM bouts. **(g)** Quantification for the number of NREM bouts after sleep onset. **(h)** Quantification of the average length of NREM bouts. **(i)** Quantification of the latency to REM sleep. **(j)** Quantification of the latency to NREM sleep. Points on **c-j** represent individual mice, n=10 mice per group. The source data and specific p-values for **c-j** are available in Figure 4-source data 1.

### Changes in delta, beta, and gamma spectral power may underlie sleep state impairments in *Ptf1a*^*Cre*^*;Vglut2*^*fx/fx*^ but not *Pdx1*^*Cre*^*;Vglut2*^*fx/fx*^ mice

Arousal states are defined, in part, by spectral frequency oscillations that occur across specific frequency bands, ranging from 0.5 to >100Hz (5A). Accordingly, changes in sleep stages are marked by changes in delta (0.5Hz – 4Hz in mice) or theta (5Hz – 8Hz in mice) power, indicating both an increase or a decrease in sleep quality.^42,43^ Therefore, changes in spectral power can give some insight as to how sleep/wake dynamics are being interrupted at the neuronal level. As dystonia is a heterogenous motor disorder^1^, and our two mouse models show differing severity in dystonic motor symptoms overall, we predicted that between the models we would see specific differences in ECoG spectral activity in sleep-dependent frequency bands. Such analysis may also provide insight into the potential mechanisms of sleep dysfunction, given that different frequency bands can be used to report on changes in overall brain connectivity^44^. We therefore sought to determine whether the *Pdx1*^*Cre*^*;Vglut2*^*fx/fx*^ and the *Ptf1a*^*Cre*^*;Vglut2*^*fx/fx*^ mice displayed measurable differences in spectral power across frequency bands of interest (Figure 5B), relative to controls. We implanted mice with two cortical (ECoG) electrodes to detect changes in oscillation power spectral frequency at various sleep stages. We calculated overall average spectral power frequency from two independent ECoG electrodes placed over the parietal cortex and the frontal cortex, to measure delta (0.5-4Hz), theta (5-8Hz), alpha (8-13Hz), beta (13-30Hz), and gamma (35-44Hz) frequency bands. *Ptf1a*^*Cre*^*;Vglut2*^*fx/fx*^, but not the *Pdx1*^*Cre*^*;Vglut2*^*fx/fx*^ mice, displayed differential spectral power frequencies in the frontal cortex for delta and beta frequency bands (Figure 5C, 5I). Specifically, delta power was significantly increased relative to the controls, while beta power was decreased. As delta power can be an effective indicator of NREM sleep, this observed increase may reflect the overall increased time spent in NREM sleep that the *Ptf1a*^*Cre*^*;Vglut2*^*fx/fx*^ mice display. We also note that although gamma power was decreased in *Ptf1a*^*Cre*^*;Vglut2*^*fx/fx*^ mice, the change did not meet the threshold for significance (Figure 5K). Overall, theta and alpha power were unchanged from controls for *Ptf1a*^*Cre*^*;Vglut2*^*fx/fx*^ and *Pdx1*^*Cre*^*;Vglut2*^*fx/fx*^ mice (Figure 5E, 5G).

**Figure 5:**
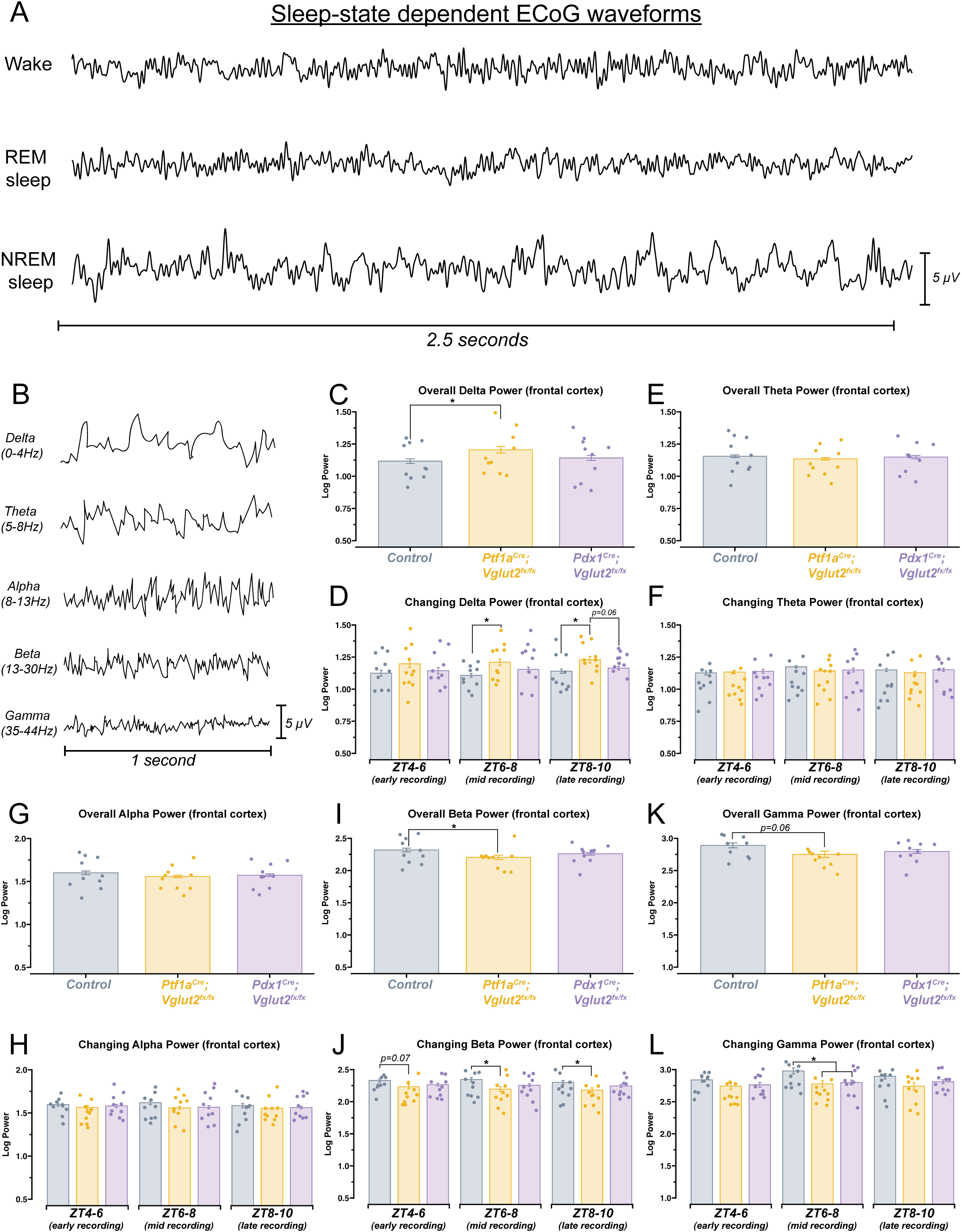
*Ptf1a*^*Cre*^*;Vglut2*^*fx/fx*^ mice show differences in spectral frequency oscillations that define arousal states. **(a)** 2.5-second samples of raw ECoG waveforms for awake, REM, and NREM from a control mouse. **(b)** 1-second samples of raw ECoG waveforms for frequency bands of interest, from a control mouse. **(c)** Quantification of delta power (0-4Hz). Average power across the entire recording period. **(d)** Quantification of the changing delta power across the recording period. **(e)** Quantification of theta power (5-8Hz). Average power across the entire recording period. **(f)** Quantification of the changing theta power across the recording period. **(g)** Quantification of alpha power (8-13Hz). Average power across the entire recording period.. **(h)** Quantification of the changing alpha power across the recording period. **(i)** Quantification of beta power (13-30Hz). Average power across the entire recording period. **(j)** Quantification of the changing beta power across the recording period. **(k)** Quantification of gamma power (35-44Hz). Average power across the entire recording period. **(l)** Quantification of the changing gamma power across the recording period. Points on **c-l** represent individual mice, n=10 mice per group. The source data and specific p-values for **c-l** are available in Figure 5-source data 1.

Sleep, and by extension the spectral oscillations defining sleep, are known to possess some intrinsic rhythmicity. Therefore, we also sought to determine whether the observed changes in spectral frequency power displayed temporal properties. To do this, we divided each animal’s sleep recording into 3 distinct periods. After performing spectral frequency analysis, we found that *Ptf1a*^*Cre*^*;Vglut2*^*fx/fx*^ mice continue to display increased delta and decreased beta power (Figure 5D, 5J). Interestingly, for both frequency bands, spectral power was only different during mid (ZT6-8) and late (ZT8-10) recording periods. For Gamma power, a significant decrease was only observed during the mid-recording period for both *Pdx1*^*Cre*^*;Vglut2*^*fx/fx*^ and *Ptf1a*^*Cre*^*;Vglut2*^*fx/fx*^ mice (Figure 5L). As we observed with overall power, neither theta nor alpha showed differences when we calculated changing power over time (Figure 5F, 5H). These data demonstrate that measurable spectral frequency changes accompany sleep impairments in dystonia, but predominantly in *Ptf1a*^*Cre*^*;Vglut2*^*fx/fx*^ mice that experience overt motor dysfunction. The data indicate the possibility that while sleep impairments arise from cerebellar dysfunction in dystonia, the overt motor defects, which can arise in parallel, can also influence specific aspects of sleep physiology in the disease.

## Discussion

We genetically dissected the interaction between sleep impairments and cerebellar-initiated motor impairments in two mouse models of dystonia. Altogether, the results from this work provide insight into the unique sleep, ECoG, and EMG disturbances observed in our mouse models of cerebellar circuit dysfunction (Figure 6A). We found that sleep impairments, a common nonmotor symptom in human dystonia, occur in *Ptf1a*^*Cre*^*;Vglut2*^*fx/fx*^ and *Pdx1*^*Cre*^*;Vglut2*^*fx/fx*^ mouse models of cerebellar miswiring, with and without severe dystonia-related motor dysfunctions. We show that both groups of mutant mice display an increase in the length of wake bouts, increased NREM and more frequent NREM bouts, and decreased REM and less frequent REM bouts (Figure 6A). While existing studies on sleep quality in dystonia patients is limited, our results are striking in that they reflect patterns of sleep deficits observed in dystonia patients^23–25^. We also highlight our finding that motor activity in *Ptf1a*^*Cre*^*;Vglut2*^*fx/fx*^ mice remains elevated in all stages of sleep, even during REM. This result is particularly intriguing. Existing studies are split, some suggesting that abnormal muscle activity in dystonic patients disappears during sleep^23^ while other indicate that it might persist during sleep^24^. On the one hand, it is possible that our results indicate that dysfunction of the mechanisms involved in synaptic renormalization are affected in dystonia, which are believed to occur during sleep and mediate muscle recovery and atonia during sleep^32,45,46^. On the other hand, as a reconciling interpretation, the work we presented here could also suggest that cerebellar dysfunction, in the presence or absence of dystonic motor dysfunction, is sufficient to drive nonmotor impairments in sleep in mouse models of dystonia (Figure 6B). Existing knowledge suggests that the cerebellum and its circuit components (namely, the Purkinje cells and cerebellar nuclei neurons) are a key node in dystonia^4,47^. Therefore, our results may point to the cerebellum as a central dystonia locus, which could help to anchor future studies on the development of therapies that can address motor and nonmotor (sleep) dysfunction in dystonia.

**Figure 6:**
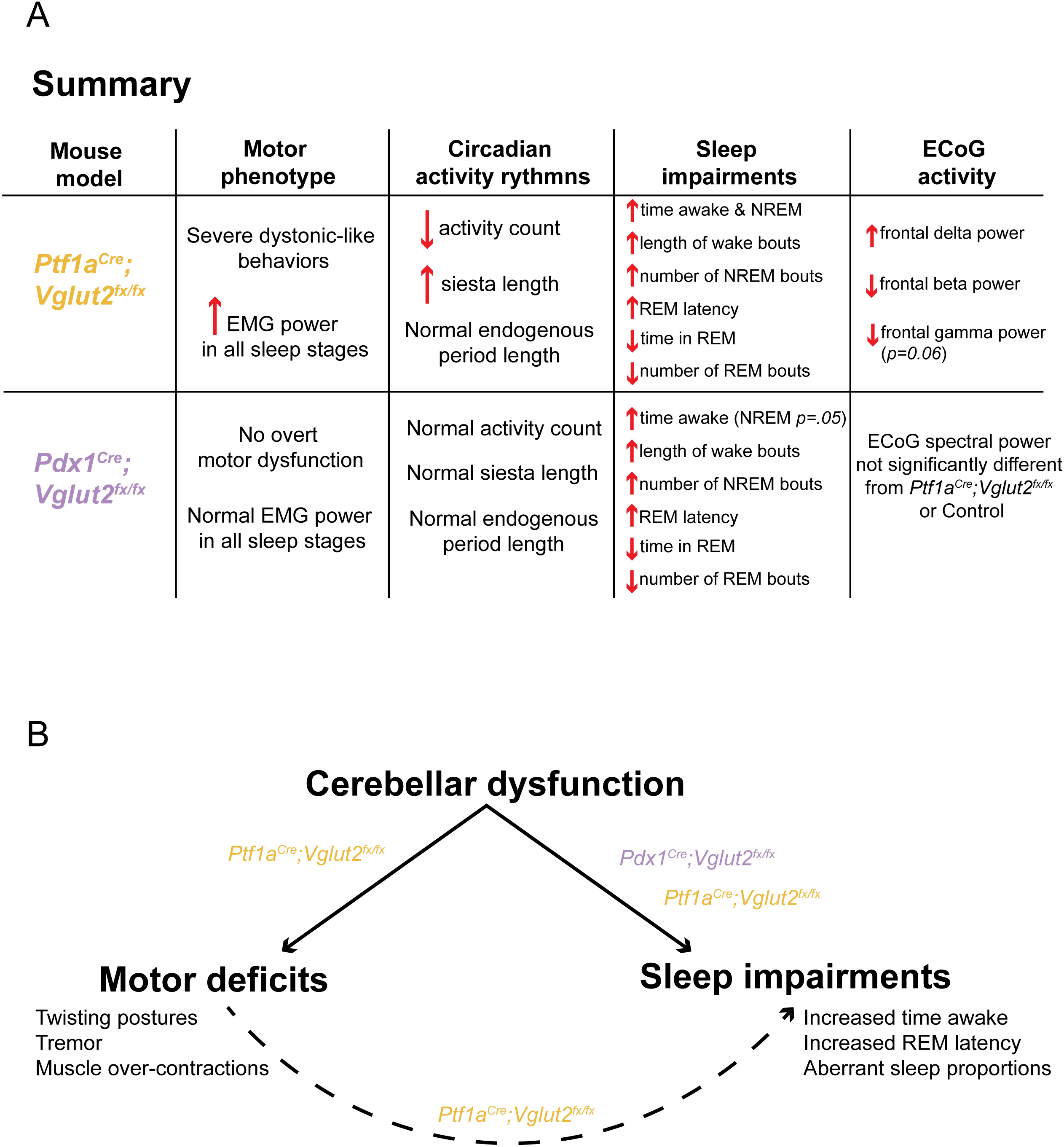
A model in which cerebellar dysfunction independently drives dystonic motor behavior and sleep impairments in *Ptf1a*^*Cre*^*;Vglut2*^*fx/fx*^ and *Pdx1*^*Cre*^*;Vglut2*^*fx/fx*^ mice. **(a)** A summary of the main findings of this study, stratified for each mouse model. **(b)** A proposed model of the main finding of this study.

Evidence from case-control studies in patients with cervical dystonia suggests that human patients with dystonia exhibit distinct abnormalities in their circadian rhythms, including fatigue and excessive daytime sleepiness^25,26,48^. We sought to understand if the clinical symptoms of dystonia that are relevant to sleep were observed in our two mouse models that exhibit cerebellar dysfunction, with and without motor deficits. Our results show that both *Pdx1*^*Cre*^*;Vglut2*^*fx/fx*^ and *Ptf1a*^*Cre*^*;Vglut2*^*fx/fx*^ mice display normal circadian timing of behavior (Figure 2C-E,H), suggesting that cerebellar dysfunction and motor deficits do not impact overall circadian behavior. Although these data are intriguing, given the substantial sleep impairments experienced by these mice, this finding is in line with work in mice with cerebellar ataxia, which also show normal circadian wheel-running behavior^37^. To make sure that our genetic manipulation in each model did not disrupt activity in the SCN master clock, we verified in our mice existing work, which states that the SCN is 95% GABAergic in its neuronal identity^39^. In accordance with our *in situ* hybridization results (Figure 2 figure supplement 1), this may indicate that while the cerebellum is involved in the regulation of sleep, its role in circadian timekeeping is limited, at least in the current context. Indeed, while numerous projections exist between the cerebellum and the major circadian centers of the brain, including the hypothalamus, locus coeruleus, and pedunculopontine nucleus, direct projections between the cerebellum and the SCN master clock are lacking^49,50^. It is possible then that cerebellar access to circadian processes is tightly regulated and restricted to sleep rather than overall activity rhythms. In this case, the fatigue and excessive daytime sleepiness experienced by patients with dystonia may be attributed to lack of sleep rather than aberrant circadian timekeeping.

While dystonia is commonly considered a network disorder in humans^1^, our genetic manipulation attempts to recreate the circuit-wide defects through a mechanism that initiates the dystonia by precisely blocking glutamatergic signaling in the cerebellum. Although both *Pdx1* and *Ptf1a* are expressed in several brain regions, including in the hypothalamus, neither *Pdx1* nor *Ptf1a* is expressed in the SCN, the circadian master clock of the brain^33,51^. Furthermore, as discussed before, over 95% of the cells in the suprachiasmatic nucleus are GABAergic^39^, which further suggests that our genetic manipulation does not extend to directly affect the master clock. Indeed, our analysis of *Vglut2* mRNA expression showed that *Vglut2* expression in the suprachiasmatic nucleus is sparse (Figure 2 figure supplement 1). As suggested by existing work, these regions instead heavily express *Vgat*, indicative of using primarily GABAergic signaling. Therefore, it is possible that human dystonia patients do experience malfunctioning circadian rhythmicity, but our model is unable to capture this specific aspect of heterogeneity in dystonia network dysfunction.

While overall circadian rhythm patterns remained unchanged in our mutant mice, we did observe a difference in siesta time for *Ptf1a*^*Cre*^*;Vglut2*^*fx/fx*^ mice. The siesta period is a brief bout of sleep during the active period^38^; it represents an important output of the circadian systems’ sleep regulation process^38,52^. Thus, it serves as an additional marker of typical circadian rhythmicity. We note that the siesta period in the *Ptf1a*^*Cre*^*;Vglut2*^*fx/fx*^ mice does appear more pronounced and is significantly longer by 7-9 minutes (Figure 2I). However, given the low overall activity profile of the *Ptf1a*^*Cre*^*;Vglut2*^*fx/fx*^ mice, it is difficult to determine whether this increase in siesta time is circadian in origin, or if it arises as a result of the decreased overall activity, making the accurate calculation of siesta onset/offset difficult. Recent work does suggest that daily timing of the siesta is under control of the SCN^53^, which we have determined to be spared from our genetic manipulation. Together, these data further support the finding that circadian rhythmicity is largely unaffected in *Ptf1a*^*Cre*^*;Vglut2*^*fx/fx*^ mice and also in the *Pdx1*^*Cre*^*;Vglut2*^*fx/fx*^ model of dystonia.

Human studies suggest that alterations in sleep efficiency and sleep latency are prevalent in dystonia patients^19,23,24,26,48,49^. We found that changes in sleep/wake dynamics are of particular interest, as they begin to explain the specific ways in which sleep deficits arise in the disease. Our results further suggest that REM disruption may be one of the primary sleep deficits encountered by our mouse models. We found not only that our mutants spend less time in REM, but that this impairment is complemented by an increase in both wake and NREM sleep (Figure 3G-I). Though mice sleep in bouts of 120-180 seconds per full sleep cycle^54^ and not in long consolidated bouts like humans, they do follow similar sleep stage patterns (Figure 3E). Despite REM representing the lightest sleep stage, it is typically preceded by NREM sleep^55^. Therefore, the increase in NREM sleep combined with decrease in REM sleep suggests that the sleep deficits in our mice specifically result from involuntary waking during NREM sleep. This is further evidenced by our EMG results for the *Ptf1a*^*Cre*^*;Vglut2*^*fx/fx*^ mice, which display elevated cervical EMG power in all sleep states (Figure 1 figure supplement 1). As mice must pass through NREM again before entering REM as they start a new sleep cycle, this prolongs the time spent in NREM while decreasing the time spent in REM. Further work needs to be conducted in order to connect our findings in mice to human patients with dystonia, as an equivalent result in humans could potentially explain the reported symptoms of daytime fatigue. Even if the total sleep time is similar between dystonic and non-dystonic patients, the quality of sleep is still being affected, as proportions of NREM versus REM during the sleeping phase are equally as important as overall time spent asleep versus awake^55^.

Given the cerebellums’ known projections to/from a variety of cortical regions involved not only in sleep regulation, but also regulation of specific sleep stages (NREM and REM)^49,50,56^, it was not unsurprising to find that sleep-stage specific deficits exist in both *Pdx1*^*Cre*^*;Vglut2*^*fx/fx*^ and *Ptf1a*^*Cre*^*;Vglut2*^*fx/fx*^ mice. Other groups have found that dystonia patients^23,26^ and mouse models of motor dysfunction^57^ present with sleep-stage specific deficits. Our findings of increased average wake bout length (Figure 4D), increased number of NREM bouts (Figure 4G) and decreased number of REM bouts (Figure 4E) specifically highlight and further reinforce our main findings of impairments in overall sleep stage timing. These results highlight the existence of significant REM-related sleep deficits. This is further reflected in our results of increased latency to reach REM for both *Pdx1*^*Cre*^*;Vglut2*^*fx/fx*^ and *Ptf1a*^*Cre*^*;Vglut2*^*fx/fx*^ mice (Figure 4I). For *Ptf1a*^*Cre*^*;Vglut2*^*fx/fx*^ mice, the motor dysfunction may partially explain this result. REM-related sleep impairments are typically accompanied by some form of motor dysfunction^24,26,57^, as the canonical mechanisms of muscle atonia during REM are disrupted^58^. However, since our *Pdx1*^*Cre*^*;Vglut2*^*fx/fx*^ mouse model does not display overt motor dysfunction, but still displays the same wake, NREM, and particularly REM-related deficits, motor dysfunction may not be the sole culprit for impaired sleep. In this case, cerebellar dysfunction may also be to blame. Indeed, the cerebellum itself and many regions receiving direct cerebellar innervation are known to be involved in sleep regulation or control sleep-dependent behaviors, particularly REM regulation^49^. The locus coeruleus regulates NREM/REM intensity^59^ while sending and receiving dense projections to cerebellar Purkinje cells and the cerebellar nuclei^60–62^. The pedunculopontine nucleus is a known regulator of REM sleep^63^ and also sends/receives inputs between cerebellum and the basal ganglia^64^. Therefore, it is possible that the cerebellar malfunctions in the *Pdx1*^*Cre*^*;Vglut2*^*fx/fx*^ and *Ptf1a*^*Cre*^*;Vglut2*^*fx/fx*^ mice directly and indirectly influence REM latency through intermediary regions such as the pedunculopontine nucleus or the locus coeruleus, or even other regions, all of which play a role in the regulation of REM sleep and receive/send direct innervation from the cerebellum^50,56,59^. The circuit pathways mediating the direct versus indirect effects on sleep were not resolved in the current work. Ultimately, the impaired sleep dynamics further reinforces our results, which suggest that *Pdx1*^*Cre*^*;Vglut2*^*fx/fx*^ and *Ptf1a*^*Cre*^*;Vglut2*^*fx/fx*^ mice experience interrupted sleep cycles, cutting REM sleep short or missing it entirely and re-starting subsequent sleep cycles from NREM sleep.

Our ECoG spectral activity results are of particular interest from a mechanistic view, as they may not only explain the factors underlying sleep deficits in *Pdx1*^*Cre*^*;Vglut2*^*fx/fx*^ and *Ptf1a*^*Cre*^*;Vglut2*^*fx/fx*^ mice but may also serve as additional “biomarkers” that differentiate each group based on their degree of cerebellar dysfunction. For instance, we observed an increase in delta power for *Ptf1a*^*Cre*^*;Vglut2*^*fx/fx*^ but not *Pdx1*^*Cre*^*;Vglut2*^*fx/fx*^ mice, which occurs predominantly in the latter stages of recording (Figure 5C, 5D). The increase for *Ptf1a*^*Cre*^*;Vglut2*^*fx/fx*^ is in agreement with recent work suggesting that higher delta power is associated with arousal and sleep impairment, particularly in the context of obstructive sleep apnea which involves both motor dysfunction and cerebellar dysfunction^65^. However, as delta power in *Pdx1*^*Cre*^*;Vglut2*^*fx/fx*^ is also elevated relative to control mice and is not significantly different from *Ptf1a*^*Cre*^*;Vglut2*^*fx/fx*^, there stands the possibility that the lack of significance stems from the “intermediate” phenotype of *Pdx1*^*Cre*^*;Vglut2*^*fx/fx*^ mice. We have shown that *Pdx1*^*Cre*^*;Vglut2*^*fx/fx*^ mice lack overt motor dysfunction (Figure 1H, 1I, Figure 1-figure supplement 2, Supplementary video 1), and that the extent of *Vglut2* deletion in the cerebellar cortex, at least with respect to the climbing fibers, is less extensive relative to the *Ptf1a*^*Cre*^*;Vglut2*^*fx/fx*^ mice (Figure 1-figure supplement 1). Therefore, if we consider each mutant group as a model for different cerebellar/motor disorders of varying intensity, we expect to see such differences in spectral power despite similar sleep deficits. In this case, spectral differences may represent “biomarkers” of disease severity. It is known that changes in delta power can differ across individuals with different diseases even if all individuals display poor sleep^43^; it is possible then that *Pdx1*^*Cre*^*;Vglut2*^*fx/fx*^ and *Ptf1a*^*Cre*^*;Vglut2*^*fx/fx*^ mice, even with an overlap in the genetic manipulations, indeed represent different manifestations of dystonic motor disease. Ultimately, this is evident with the difference in observed dystonic motor phenotypes between the two groups. It should be noted though, the measurement of sleep-related spectral difference between the *Pdx1*^*Cre*^*;Vglut2*^*fx/fx*^ and *Ptf1a*^*Cre*^*;Vglut2*^*fx/fx*^ mice could still intersect with the induced alterations in the motor program, as movement patterns are known to impact ECoG spectral activity^66^.

Additional patterns of significantly different spectral power for *Ptf1a*^*Cre*^*;Vglut2*^*fx/fx*^ but not *Pdx1*^*Cre*^*;Vglut2*^*fx/fx*^ mice are seen for beta power, indicative of alert wakefulness (Figure 5I, 5J). While beta power is typically increased in patients with primary insomnia, previous work has shown that decreased beta activity is also associated with poor sleep quality, particularly in patients with obstructive sleep apneas^65^. The relationship between the cerebellum and breathing is well-established and may provide a fruitful avenue for further research in the context of dystonia. The cerebellum is known to be involved in both the rhythmicity of breathing and in regulating air hunger^67^; both mechanisms are known to play a role in obstructive sleep apneas^19,68^. It is possible that *Ptf1a*^*Cre*^*;Vglut2*^*fx/fx*^ mice, with overt motor dysfunction, have some degree of sleep apnea behavior, which could contribute to their observed sleep impairment. Interestingly, cortical gamma power was only significantly changed during the middle of the recording period, for both the *Pdx1*^*Cre*^*;Vglut2*^*fx/fx*^ and the *Ptf1a*^*Cre*^*;Vglut2*^*fx/fx*^ mutant mice (Figure 5L). While gamma oscillations are typically associated with working memory and attention, human and mouse EEG/ECoG studies have found that gamma oscillations occur spontaneously during REM and NREM sleep^69,70^. This may explain why the overall gamma power is not significantly lower for either the *Pdx1*^*Cre*^*;Vglut2*^*fx/fx*^ or *Ptf1a*^*Cre*^*;Vglut2*^*fx/fx*^ mice, yet it does reach the threshold for significance during “mid-recording”. *Ptf1a*^*Cre*^*;Vglut2*^*fx/fx*^ and *Pdx1*^*Cre*^*;Vglut2*^*fx/fx*^ mice exhibit an increase in NREM and a decrease in REM sleep, and gamma oscillations spontaneously occur during both stages; changes in gamma activity may be effectively “canceled out” due to the opposing NREM and REM dynamics. While these observed changes in spectral frequency oscillations uncover some of the potential mechanisms driving the observed changes in sleep in our *Ptf1a*^*Cre*^*;Vglut2*^*fx/fx*^ and *Pdx1*^*Cre*^*;Vglut2*^*fx/fx*^ mice, we note that attributing sleep disruption to specific directional changes in any one frequency band is difficult. Both positive and negative changes in average power, in any frequency band, can be associated with various disease states, and notably with disordered sleep^42,43^. Therefore, here, we highlight the presence of a change in spectral frequency power as an indicator of fractured sleep homeostasis in our mouse circuit models of dystonia, without differentiating the specific directionality of the change in spectral frequency power.

Our findings build upon existing evidence from both human patients and mouse models of motor disease demonstrating that sleep impairments are a common nonmotor symptom in dystonia. Previous work has been unable to distinguish between dystonia-dependent versus independent sleep dysfunction, particularly in the context of dystonic motor dysfunction. Importantly, our results suggest a model in which cerebellar dysfunction alone (Figure 6B), without overt dystonic motor phenotypes, can drive sleep deficits. This may be an indication of a broader set of network dysfunctions in dystonia, with the cerebellum located at the center of multiple disease symptoms.

## Ethics

Animal experimentation: All animals were housed in an AALAS-certified facility that operates on a 14 hour light cycle. Husbandry, housing, euthanasia, and experimental guidelines were reviewed and approved by the Institutional Animal Care and Use Committee (IACUC) of Baylor College of Medicine (protocol number: AN-5996).

## Acknowledgements and Funding Sources

This work was supported by Baylor College of Medicine (BCM), Texas Children’s Hospital, The Hamill Foundation, and the National Institutes of Neurological Disorders and Stroke (NINDS) R01NS100874, R01NS119301, and R01NS127435 to RVS. Research reported in this publication was supported by the Eunice Kennedy Shriver National Institute of Child Health & Human Development of the National Institutes of Health under Award Number P50HD103555 for use of the Animal Behavior Core and the Cell and Tissue Pathogenesis Core (BCM IDDRC). The content is solely the responsibility of the authors and does not necessarily represent the official views of the National Institutes of Health. Support was also provided by a Dystonia Medical Research Foundation (DMRF) grant to RVS.

Research reported in this publication was supported in part by the RNA *In Situ* Hybridization Core facility at Baylor College of Medicine, which is supported by a Shared Instrumentation grant from the NIH (S10OD016167) and the NIH IDDRC grant U54HD083092 from the Eunice Kennedy Shriver National Institute of Child Health & Human Development. The content is solely the responsibility of the authors and does not necessarily represent the official views of the Eunice Kennedy Shriver National Institute of Child Health & Human Development or the National Institutes of Health.

## Author contributions

Technical and conceptual ideas in this work were conceived by LESL and RVS. LESL performed the experiments and LESL and RVS performed data analysis and data interpretation. LESL and RVS wrote and edited the manuscript.

## Conflicts of interest

We have no conflicts of interest to disclose.

## Figure Legends

**Figure 1-figure supplement 1:**
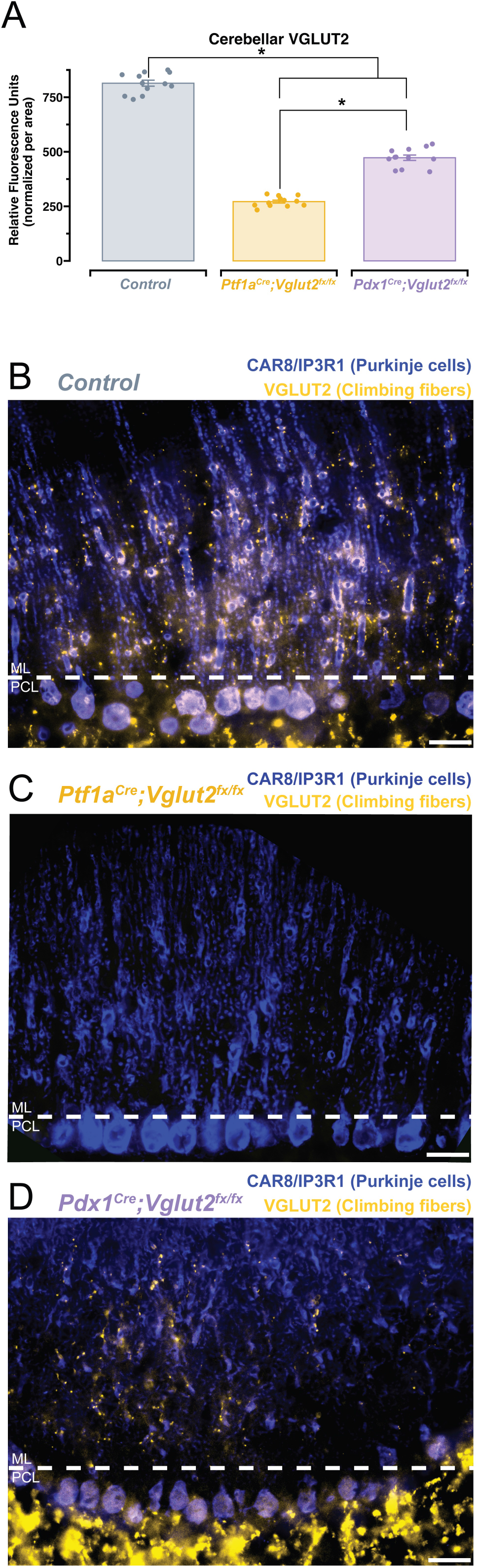
Deletion of VGLUT2 in the cerebellum is more widespread in the climbing fibers of the *Ptf1a*^*Cre*^*;Vglut2*^*fx/fx*^ mice than in the *Pdx1*^*Cre*^*;Vglut2*^*fx/fx*^ mice. **(a)** Quantification of relative fluorescence units (normalized per area) for each group. **(b)** Fluorescence immunohistochemical stain of the cerebellar cortex for a control mouse. Purkinje cell bodies and axons are shown in blue (labeled with CAR8/IP3R1). Climbing fibers express VGLUT2 and are labeled in gold. Scale bars are 20um and indicated with white bars. The molecular layer (ML) and Purkinje cell layer (PCL) are labeled for orientation. **(c)** Same as **(b)** but for a *Ptf1a*^*Cre*^*;Vglut2*^*fx/fx*^ mouse. **(d)** Same as **(b)** but for a *Pdx1*^*Cre*^*;Vglut2*^*fx/fx*^ mouse. Points on **a** represent individual sections (n=4) from 3 mice per group. The source data and specific p-values for **a** are available in Figure 1-source data 1.

**Figure 1-figure supplement 2:**
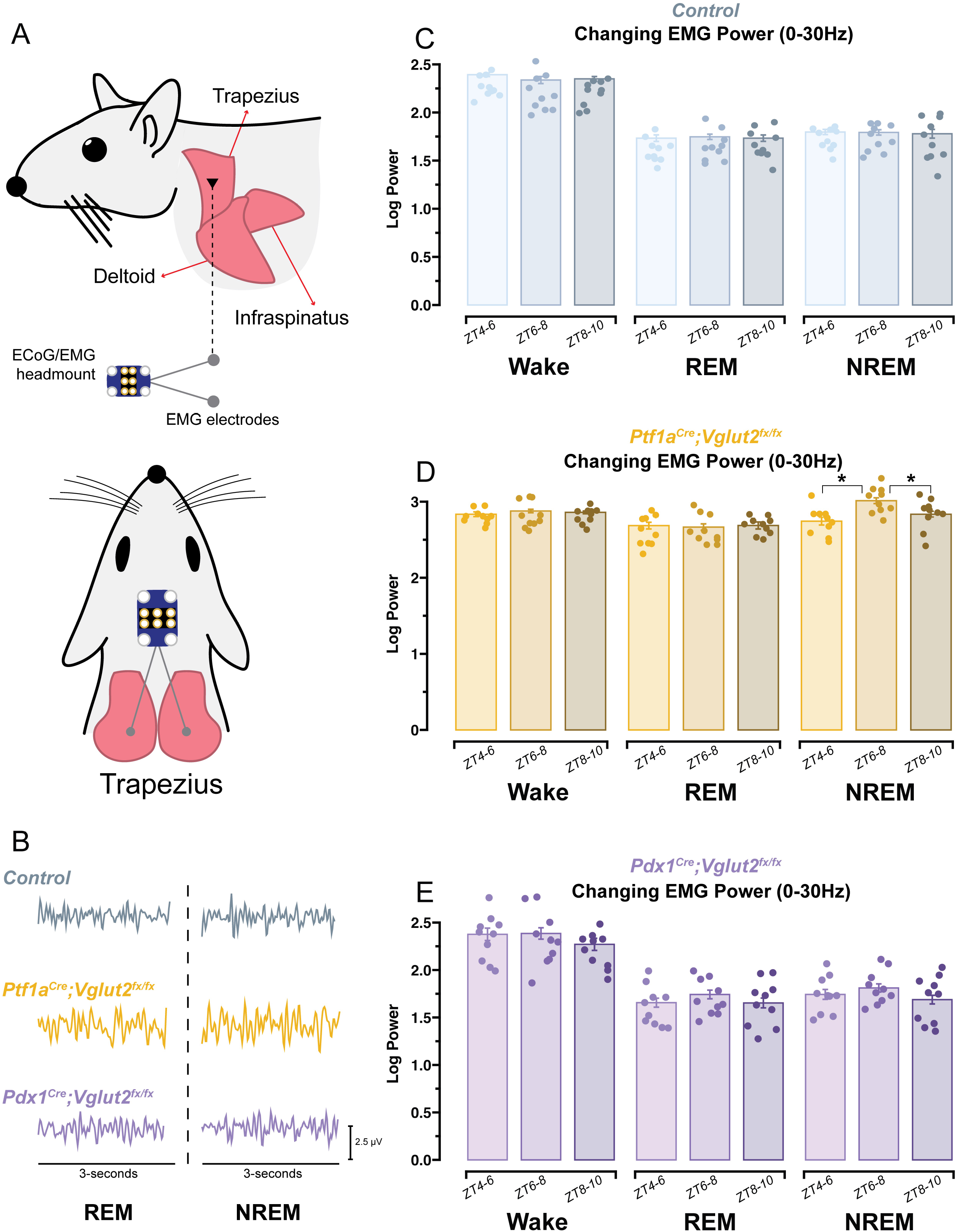
Cervical EMG activity in *Ptf1a*^*Cre*^*;Vglut2*^*fx/fx*^ mice remains elevated in all states. **(a)** Schematic illustration of a mouse showing the musculature and relative placement of the EMG electrodes. **(b)** Raw EMG waveforms of trapezius activity (3-seconds) for REM and NREM sleep for mice of each group. **(c)** Quantification of changing EMG power for control mice. ZT0 = lights ON, ZT4 = 4-hr after lights on… etc. **(d)** Same as **(c)** but for *Ptf1a*^*Cre*^*;Vglut2*^*fx/fx*^ mice. **(e)** Same as **(c)** but for *Pdx1*^*Cre*^*;Vglut2*^*fx/fx*^ mice. Points on **c-e** represent individual mice, n=10 mice per group. The source data and specific p-values for **c-e** are available in Figure 1-source data 1.

**Figure 2-figure supplement 1:**
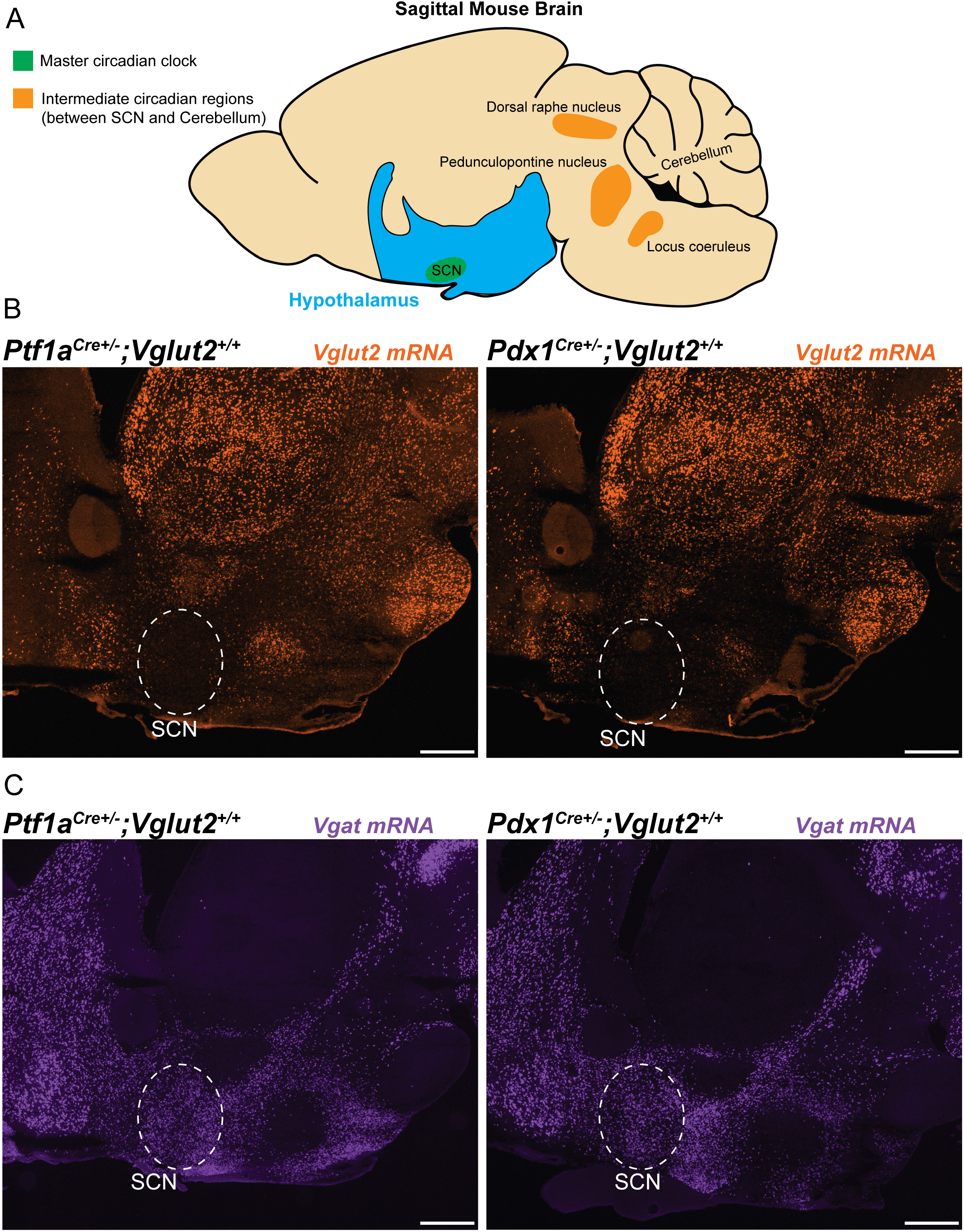
The major circadian centers of the hypothalamus are comprised primarily of GABAergic inhibitory neurons. **(a)** Schematic illustration of a sagittal mouse brain highlighting several brain regions of interest that are related to circadian behavior. The hypothalamus is shown in blue, the suprachiasmatic nucleus (SCN) is in green, and the intermediate regions located between the SCN-cerebellum are shown in orange. **(b)** Images processed using *in situ* hybridization revealing *Vglut2* mRNA expression on sagittal sections cut through the adult mouse brain. The regions of interest are outlined in white. **(c)** Same as **(b)** but for *Vgat* mRNA expression.

**Supplementary Video 1: Differential motor phenotypes in *Pdx1*^*Cre*^*;Vglut2*^*fx/fx*^ and *Ptf1a*^*Cre*^*;Vglut2*^*fx/fx*^ mice.**

A single video of the mutant mice showing the absence (*Pdx1*^*Cre*^*;Vglut2*^*fx/fx*^) and presence (*Ptf1a*^*Cre*^*;Vglut2*^*fx/fx*^) of dystonic motor behaviors.

## Methods

### Animals

All mice used in this study were housed in a Level 3, AALAS-certified facility. All experiments and studies that involved mice were reviewed and approved by the Institutional Animal Care and Use Committee of Baylor College of Medicine (BCM AN-5996). Dr Chris Wright (Vanderbilt University School of Medicine) kindly provided the *Ptf1a*^*Cre*^ mice. We purchased the *Pdx1*^*Cre*^ (Pdx-Cre, #014647) and *Vglut2*^*floxed*^ (*Vglut2*^*fx*^, #012898) mice from The Jackson Laboratory (Bar Harbor, ME, USA) and then maintained them in our colony using a standard breeding scheme. The conditional knock-out mice that resulted in dystonia were generated by crossing *Ptf1a*^*Cre*^*;Vglut2*^*fx/fx*^ heterozygote mice or *Pdx1*^*Cre*^*;Vglut2*^*fx/fx*^ heterozygote mice with homozygote *Vglut2*^*fx/fx*^ mice. *Pdx1*^*Cre*^*;Vglut2*^*fx/fx*^ and *Ptf1a*^*Cre*^*;Vglut2*^*fx/fx*^ mice were considered experimental animals. A full description of the genotyping details (e.g., primer sequences and the use of a standard polymerase chain reaction) and phenotype for the *Ptf1a*^*Cre*^*;Vglut2*^*fx/fx*^ mouse was provided in White and Sillitoe, 2017^4^. A full description of the genotype and the initial observations of the phenotype of the *Pdx1*^*Cre*^*;Vglut2*^*fx/fx*^ mouse was provided in Lackey, 2022^34^. All littermates lacking Cre upon genotyping were considered control mice. Ear punches were collected before weaning and used for genotyping and identification of the different alleles. For all experiments, we bred mice using standard timed pregnancies, noon on the day a vaginal plug was detected was considered embryonic day (E)0.5 and postnatal day (P)0 was defined as the day of birth. Mice of both sexes were used in all experiments.

### Immunohistochemistry

Perfusion and tissue fixation were performed as previously described^71^. Briefly, mice were anesthetized by intraperitoneal injection with Avertin (2, 2, 2-Tribromoethanol, Sigma-Aldrich, St. Louis, MO, USA; catalog #T4). Cardiac perfusion was performed with 0.1 M phosphate-buffered saline (PBS; pH 7.4), then by 4% paraformaldehyde (4% PFA) diluted in PBS. For cryoembedding, brains were post-fixed at 4 °C for 24 to 48 h in 4% PFA and then cryoprotected stepwise in sucrose solutions (15% and 30% diluted in PBS) and embedded in Tissue-Tek O.C.T. compound (Sakura Finetek, Torrance, CA, USA; catalog #4583). Tissue sections were cut on a cryostat at a thickness of 40 μm and individual free-floating sections were collected sequentially and immediately placed into PBS. Our procedures for immunohistochemistry on free-floating frozen cut tissue sections have been described extensively in previous work^72,73^. After completing the staining steps, the tissue sections were placed on electrostatically coated glass slides and allowed to dry.

Rabbit polyclonal anti-CA VIII (CAR8, 1:500, Proteintech # 12391-1-AP) and rabbit polyclonal anti-IP3R1 (1:500, Invitrogen # PA1-901) were used to label Purkinje cells. Guinea pig polyclonal anti-VGLUT2 (1:500, Synaptic systems # 135 404) was used to label olivocerebellar climbing fibers and their terminals. We visualized immunoreactive complexes using anti-rabbit or anti-guinea pig secondary antibodies conjugated to Alexa-488 and -647 fluorophores (1:1000 for both, Invitrogen, Waltham, MA, USA).

### Tissue preparation and processing for *in situ* hybridization

Mice were anaesthetized with isoflurane and brains were removed from the skull and immersed in OCT (optimal cutting temperature). Immersed brains were flash frozen by placing tissue molds onto dry ice. Sagittal sections (25 μm) were cut through the cerebellum and the slices placed onto electrostatically coated glass slides (Probe On Plus Fisher Brand; Fisher Scientific). The tissue was probed with *Vglut2* (*SLC17A6*) or *Vgat* (*SLC32A1*) digoxigenin-labelled mRNA probes using an automated *in situ* hybridization procedure (Genepaint). All reagent incubations, washes and stains were automated and performed by the *in situ* hybridization robot. The signal was detected by colorimetric detection using BCPI/NBT reagents. After processing was completed, the slides were removed from the machine and then cover-slipped with permanent mounting medium (Entellan mounting media, Electron Microscopy Sciences, Hatfield, PA, USA) and left to dry before imaging.

### Wheel-running behavior

Recordings were maintained in a ventilated, temperature-controlled, and light-tight room under either a 12:12 LD cycle or DD conditions. Mice were singly housed in wheel-running cages and allowed to entrain to the LD cycle for 2-weeks, before being released into DD conditions for 21-days, to assess endogenous circadian timekeeping ability. We assessed period length, activity onset, and average number of wheel revolutions per 5-minutes using ClockLab Analysis (Actimetrics).

### ECoG/EMG sleep recordings

Mice were anesthetized with isoflurane and placed into a stereotaxic device, which continued to deliver isoflurane throughout surgery. Each mouse with implanted with a prefabricated ECoG/EMG headmount (Pinnacle Technology, Lawrence KS, #8201) with 0.10” EEG screws to secure headmounts to the skull (Pinnacle Technology, Lawrence KS, #8209). A midline incision was made, and the skull was exposed. The headmount was affixed to the skull using cyanoacrylate glue to hold in place while pilot holes for screws were made and screws were inserted. Screws were placed bilaterally over parietal cortex and frontal cortex. A small amount of silver epoxy (Pinnacle Technology, Lawrence KS, #8226) was applied to the screw-headmount connection. Platinum-iridium EMG wires on the prefabricated headmount were placed under the skin of the neck, resting directly on the trapezius muscles. The headmount was permanently affixed to the skull using ‘Cold-Cure’ dental cement (A-M systems, #525000 and #526000). Mice were allowed to recover for 3-4 days before being fitted with a preamplifier (#8202) and tethered to the recording device (#8204 and #8206-HR). ECoG and EMG signals were sampled at 400Hz with 0.5Hz and 10Hz high-pass filters respectively.

Mice were recorded in light and temperature-controlled rooms, for 8-hours, at the same time of day for every mouse. The first hour of recording was considered the acclimation period and was therefore excluded from final analysis. Food and water were available *ad libitum* throughout the recording day.

### Sleep scoring and analysis of sleep data

Sleep was automatically scored offline via SPINDLE^74^. For spectral frequency analysis of ECoG and EMG activity, raw files were also pre-processed in MATLAB (MathWorks) using the free toolkit EEGLAB (UC San Diego). Scored files were downloaded from SPINDLE as a .csv and statistical analysis was performed in R v4.1.2. Only ECoG spectral power from frontal cortex is discussed in depth, as the spectral power from parietal cortex was the same between all groups for all frequency bands: Delta: one-way ANOVA, F = 0.031, p = 0.97, Theta: one-way ANOVA, F = 0.663, p = 0.524, Alpha: one-way ANOVA, F = 0.327, p = 0.724, Beta: one-way ANOVA, F = 0.079, p = 0.924, Gamma: one-way ANOVA, F = 0.483, p = 0.622.

## Data analysis and statistics

Data are presented as mean ± SEM and analyzed as a one-way ANOVA followed by Tukey’s Honest Significant Difference test for post-hoc comparisons or a repeated measures two-way ANOVA with Bonferroni correction for multiple comparisons. p < 0.05 was considered as statistically significant. All statistical analyses were performed using R v4.1.2.

## Data availability

All data generated or analyzed in this study are included in the manuscript and supporting files.

## Notes

### Competing Interest Statement

The authors have declared no competing interest.

